# Anti-COVID-19 efficacy of ivermectin in the golden hamster

**DOI:** 10.1101/2020.11.21.392639

**Authors:** Guilherme Dias de Melo, Françoise Lazarini, Florence Larrous, Lena Feige, Lauriane Kergoat, Agnès Marchio, Pascal Pineau, Marc Lecuit, Pierre-Marie Lledo, Jean-Pierre Changeux, Hervé Bourhy

## Abstract

The devastating coronavirus disease 2019 (COVID-19) pandemic, due to SARS-CoV-2, has caused more than 47 million confirmed cases and more than 1.2 million human deaths around the globe^1^, and most of the severe cases of COVID-19 in humans are associated with neurological symptoms such as anosmia and ageusia, and uncontrolled inflammatory immune response^2–5^. Among therapeutic options^6–8^, the use of the anti-parasitic drug ivermectin (IVM), has been proposed, given its possible anti-SARS-CoV-2 activity^9^. Ivermectin is a positive allosteric modulator of the α-7 nicotinic acetylcholine receptor^10^, which has been suggested to represent a target for the control of Covid-19 infection^11^, with a potential immunomodulatory activity^12^. We assessed the effects of IVM in SARS-CoV-2-intranasally-inoculated golden Syrian hamsters. Even though ivermectin had no effect on viral load, SARS-Cov-2-associated pathology was greatly attenuated. IVM had a sex-dependent and compartmentalized immunomodulatory effect, preventing clinical deterioration and reducing olfactory deficit in infected animals. Importantly, ivermectin dramatically reduced the *Il-6/Il-10* ratio in lung tissue, which likely accounts for the more favorable clinical presentation in treated animals. Our data support IVM as a promising anti-COVID-19 drug candidate.

---

Coronaviruses cause respiratory disease in a wide variety of hosts. During the ongoing pandemic of COVID-19 caused by SARS-CoV-2, clinical signs other than respiratory symptoms have been linked to infection, frequently associated with an altered sense of smell and by high mortality in some COVID-19 patients. These features seem related to the over-responsiveness of patients’ immune system to SARS-CoV-2, sometimes referred to as ‘cytokine storm’^4,5,13^.

Ivermectin (IVM), a macrocyclic lactone, is a commercially-available anti-parasitic drug which induces effects in various endo- and ectoparasites, mycobacteria and even some viruses^12,14^. IVM is an efficient positive allosteric modulator of the α-7 nicotinic acetylcholine receptor (nAChR) and of several ligand-gated ion channels, including the muscle receptor for glutamate (GluCl) in worms^10^. Furthermore, IVM has been shown to modulate the host’s immune system^12,14^ under conditions that are known to involve the α-7 receptor^15^. An IVM anti-COVID-19 effect has been hypothesized due to similarities between SARS-CoV-2 and other nicotinic receptor ligand sequences^11^. *In vitro* IVM inhibition of SARS-CoV-2 replication in Vero/hSLAM cells^9^ has also been reported, albeit at much higher concentrations (50-to 100-fold) than those clinically attainable in human patients (150-400 μg/kg)^16–18^.

Several clinical trials using IVM have been initiated either alone or in combination with other molecules^19^. IVM has already been administered to hospitalized COVID-19 patients, with contrasting outcomes: one study related no efficacy of late IVM administration (8-18 days after symptoms onset) in severe COVID-19 patients treated in combination with other drugs (hydroxychloroquine, azithromycin, tocilizumab, steroids)^20^, whereas another study observed lower mortality, especially in severe C0VID-19 patients treated with IVM in addition to usual clinical care (hydroxychloroquine, azithromycin, or both)^21^. Finally, in a pilot clinical trial, Gorial *et al.* reported a reduction in the hospital stay for the patients receiving IVM on admission, as add-on therapy with hydroxychloroquine/azithromycin^22^.

Consequently, the aim of this study was to investigate the effects of IVM alone on SARS-CoV-2 infection using the golden Syrian hamster as a model for COVID-19^23^. Male and female adult golden Syrian hamsters were intranasally inoculated with 6×10^4^ PFU of SARS-CoV-2 [BetaCoV/France/IDF00372/2020]. This inoculum size was selected as it invariably causes symptomatic infection in golden Syrian hamster, with a high incidence of anosmia and high viral loads in the upper and lower respiratory tracts within four days post-infection^24^. At the time of infection, animals received a single subcutaneous injection of IVM at the anti-parasitic dose of 400 μg/kg classically used in a clinical setting and were monitored over four days. Mock-infected animals received the physiological solution only.

IVM-treated and infected animals exhibited a significant reduction in the severity of clinical signs (Fig. 1a) and remarkably, IVM treatment reduced the olfactory deficit in infected animals: 66.7% (12/18) of the saline-treated hamsters presented with hyposmia/anosmia, whereas only 22.2% (4/18) of IVM-treated hamsters presented signs of olfactory dysfunction (Fisher’s exact test p=0.018; Fig. 1a, Extended Data Fig. 1). This effect was sex-dependent: infected males presented a reduction in the clinical score (Fig. 1b) whereas a complete absence of signs was noticed in the infected females (Fig. 1c). Regarding the olfactory performance, 83.3% (10/12) of the saline-treated males presented with hyposmia/anosmia, in contrast to only 33.3% (4/12) of IVM-treated males (Fisher’s exact test p=0.036). Furthermore, no olfactory deficit was observed in IVM-treated females (0/6), while 33.3% (2/6) of saline-treated females presented with hyposmia/anosmia (Fisher’s exact test p=0.455). The IVM-treated and infected animals presented, however, a transient decrease of body weight similar to that observed in saline-treated and infected hamsters (Extended Data Fig. 1).

**Fig. 1.**
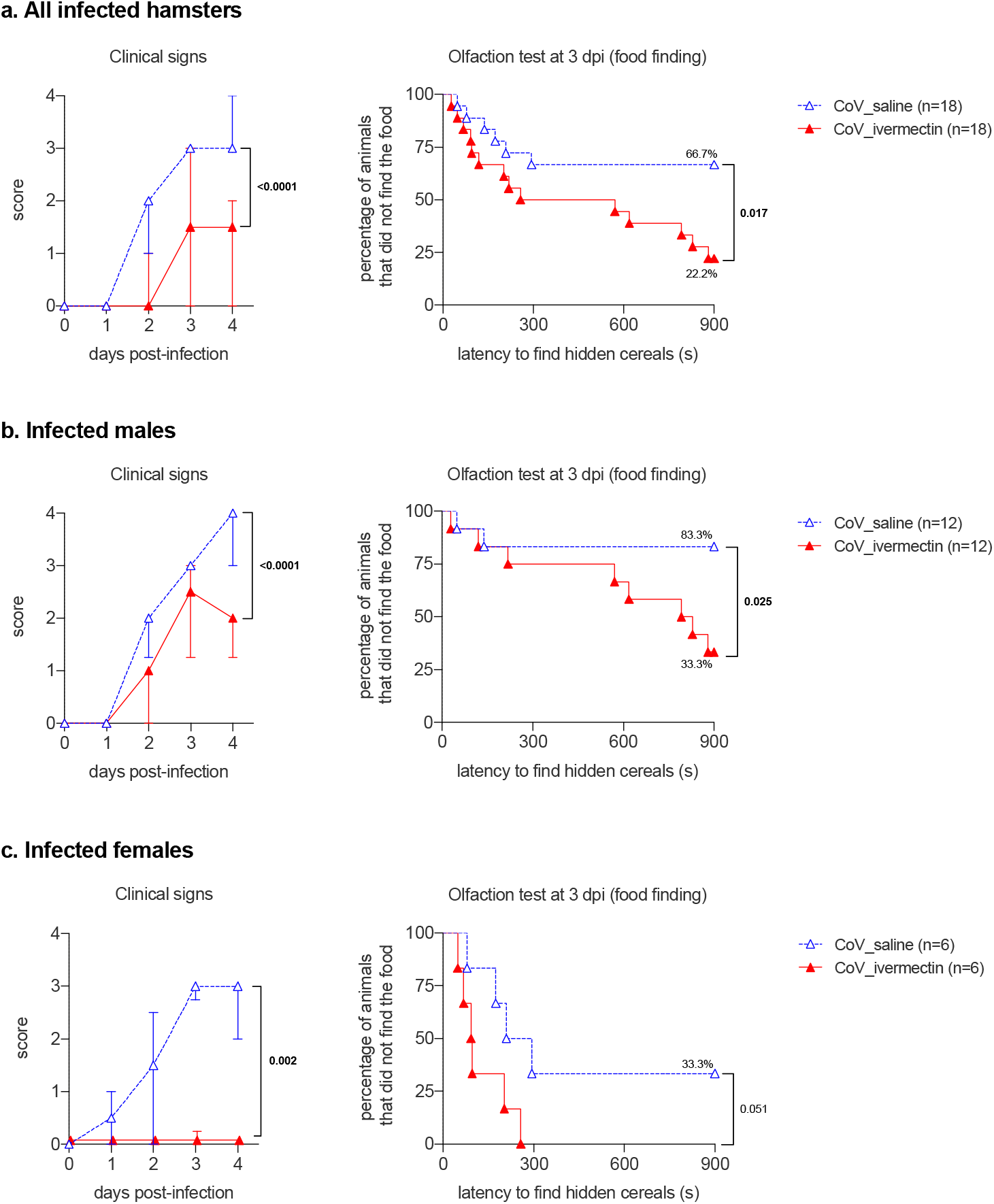
Clinical presentation and olfaction test of SARS-CoV-2-infected hamsters with and without ivermectin treatment. **a.** clinical signs and olfactory deficit in all infected hamsters. **b,** clinical signs and olfactory deficit in infected male hamsters only. **c,** clinical signs and olfactory deficit in infected female hamsters only. The clinical score is based on a cumulative 0-4 scale: ruffled fur; slow movements; apathy; stress when manipulated. The olfaction test is based on the buried food finding test. Curves represent the percentage of animals that did not find the buried food. Food finding assays were performed at 3 days post-infection. Mann-Whitney test at 4 dpi (clinical signs) and Logrank (Mantel-Cox) test (olfaction tests). The p value is indicated in bold when significant at a 0.05 threshold. Symbols indicate the median ± interquartile range. Data were obtained from three independent experiments for males and two independent experiments for females. See extended Fig. 1.

Since males presented a high index of anosmia/hyposmia, we subsequently performed a dose-response curve to test the effect of IVM on the clinical presentation of infected males: lower doses of IVM (100 or 200 μg/kg) elicited similar clinical outcomes as the anti-parasitic dose of 400 μg/kg (Extended Data Fig. 2). As expected, no signs of olfactory deficit were observed in the mock-infected hamsters (Extended Data Fig. 1). Despite sex differences, IVM treatment reduced clinical deterioration in SARS-CoV-2-infected hamsters, and even prevented the occurrence of anosmia, a typical symptom of COVID-19 in humans^5,13^.

A panel of selected cytokines (*Il-6, Il-10, Il-1β, Tnf-α* and *Ifn-γ*) and chemokines (*Cxcl10* and *Ccl5*), already known to be affected in COVID-19^4^ disease progression in humans and animal models^23^, were used to assess the impact of IVM treatment on the immune response of SARS-CoV-2 infected hamsters. We assayed two airway compartments: nasal turbinates and lungs.

In the nasal turbinates, upon treatment with IVM, there were marked differences between sex groups: females presented an important down-regulation of several mediators (*Il-6, Il-10, Tnf-α* and *Cxcl10*) while males presented an increase in two pro-inflammatory mediators (*Ifn-γ* and *Ccl5*) (Fig. 2a, Extended Data Fig. 3). Further, the expression of *Cxcl10*^25^–a key mediator known to be involved in respiratory disease and olfaction dysfunction in COVID-19 patients–, was remarkably lower in the nasal turbinates of IVM-treated females without significant changes in males. These findings are in line with the better performance of IVM-treated females observed in the food finding tests (Fig. 1).

**Fig. 2.**
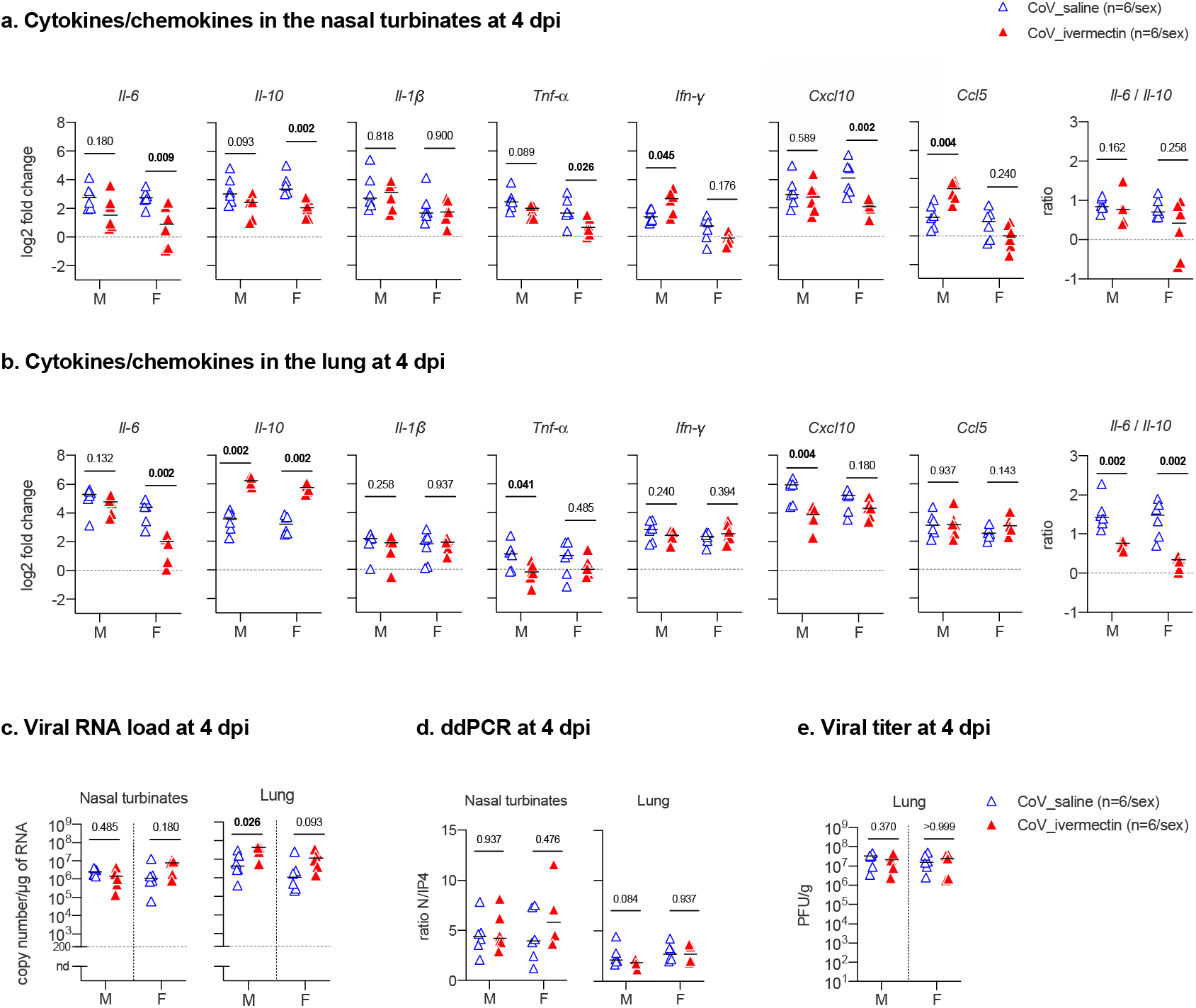
Immunological and virologic aspects in the nasal turbinates and in the lungs at 4 days-post-infection of SARS-CoV-2-infected hamsters with and without ivermectin treatment. **ab**, cytokines and chemokines transcripts in the nasal turbinates (a) and in the lungs (b) at 4dpi in male and female SARS-CoV-2 infected hamsters, treated with saline or with 400 μg/kg ivermectin. **c**, viral load in the nasal turbinates and in the lungs at 4 dpi. **d**, ratio between the CPD (copy per droplets, normalized to *γ-actin* and *Hprt* reference genes relative expression) of structural [N, nucleocapsid] and non-structural [IP4: RdRp, RNA-dependent RNA polymerase] viral gene expression determined by digital droplet PCR (ddPCR) in the nasal turbinates and in the lungs at 4 dpi. **e**, infectious viral titer in the lung at 4 dpi expressed as Plaque Forming Units (PFU)/g of tissue. Mann-Whitney test. The p value is indicated in bold when significant at a 0.05 threshold. Horizontal lines indicate medians. M: male hamsters; F: female hamsters. Data were obtained from two independent experiments for each sex. See extended Fig. 3–5.

In the lungs, however, the significant overexpression of *Il-10* was a common feature of IVM-treated males and females (Fig. 2b, Extended Data Fig. 4). This effect may be related to a modulation of the inflammatory response in the lung (down-regulation of *Tnf-α* and *Cxcl10* in males, and of *IL-6* in females) associated with the reduced clinical signs. Additionally, the *Il-6/Il-10* ratio in the lung of IVM-treated hamsters was significantly lower than in non-treated animals (Fig. 2b).

The modulation of some immune mediators in the IVM-treated hamsters confirms what was observed in other infectious contexts and in other animal species^12,14^. In particular, the low *Il-6/Il-10* ratio observed in the lung of IVM-treated hamsters may predict their better clinical presentation, as observed in humans, as lower plasmatic *IL-6/IL-10* ratios are detected in hospitalized COVID-19 patients who do not require intensive care^26^.

The viral RNA load in the respiratory tract remained unaffected by IVM treatment in both nasal turbinates and lung samples. These were tested using both classical RT-qPCR (Fig. 2c) and the highly sensitive technique of digital droplet PCR^27^ (Fig. 2d, Extended Data Fig. 5). Furthermore, IVM treatment did not influence the viral replication rate, as evaluated by the ratio between structural and non-structural gene transcription (Fig. 2d, Extended Data Fig. 5). Finally, IVM treatment did not alter infectious viral titers in the lungs (Fig. 2e). These results differ from a previous report suggesting that IVM, albeit used at far higher concentrations, inhibits the replication of SARS-CoV-2 *in vitro*^9^. Therefore, the action of IVM on COVID-19 signs in the golden hamster model does not result from its antiviral activity.

In humans, IVM is widely used as anti-helminthic and anti-scabies at therapeutic doses (150-400 μg/kg)^16^ that are in the range of those used in our golden hamster experiments. Moreover, considerable sex differences are observed in terms of clinical presentation in hamsters, as is seen in human COVID-19 patients, where men tend to develop more severe disease than women^28,29^. Further studies are needed to assess the effects of IVM when administered at later time-points during the course of the disease. Considering the results observed in our golden hamster model, together with their limitations, IVM may be considered as a new category of therapeutic agent against COVID-19 which would not modify SARS-CoV-2 replication but affect the pathophysiological consequences of the virus *in vivo*. Indeed, a characteristic modulation of the cytokine gene expression in the airways was observed in IVM-treated hamsters that led to a cytokine profile similar with that observed in humans exhibiting milder symptoms and presenting improved prognosis^26^.

The molecular mechanism of IVM’s anti-COVID-19 effect remains largely unknown. Even if the effects observed may, to some extent, share similarities with the anti-inflammatory action of dexamethasone^7^ and tocilizumab (anti-IL-6)^8^. IVM effects are steady and strong in the golden hamsters and thought to occur upstream of inflammatory pathways. Among possible molecular targets, nAChRs have been suggested to directly or indirectly interact with SARS-CoV-2^11^. An interaction of IVM with the α-7 receptor^10^ may account for the modulation of the immune response in treated hamsters, an interpretation coherent with the mobilization of the cholinergic antiinflammatory pathway (CAP) under vagus nerve control^15^, as observed in other models^30^. On the basis of these findings, and given its innocuity, we conclude that IVM should be considered as a promising anti-COVID-19 drug candidate, alone or as add-on therapy.

## Methods

### Ethics

All animal experiments were performed according to the French legislation and in compliance with the European Communities Council Directives (2010/63/UE, French Law 2013–118, February 6, 2013) and according to the regulations of Pasteur Institute Animal Care Committees. The Animal Experimentation Ethics Committee (CETEA 89) of the Institut Pasteur approved this study (200023; APAFIS#25326-2020050617114340 v2) before experiments were initiated. Hamsters were housed by groups of 4 animals in isolators and manipulated in class III safety cabinets in the Pasteur Institute animal facilities accredited by the French Ministry of Agriculture for performing experiments on live rodents. All animals were handled in strict accordance with good animal practice.

### Production and titration of SARS-CoV-2 virus

The isolate BetaCoV/France/IDF00372/2020 (EVAg collection, Ref-SKU: 014V-03890) was kindly provided by Sylvie Van der Werf. Viral stocks were produced on Vero-E6 cells infected at a multiplicity of infection of 1×10^-4^ PFU (plaque-forming units). The virus was harvested 3 days post-infection, clarified and then aliquoted before storage at −80°C. Viral stocks were titrated on Vero-E6 cells by classical plaque assays using semisolid overlays (Avicel, RC581-NFDR080I, DuPont)^31^.

### SARS-CoV-2 model and ivermectin treatment of hamsters

Male and female Syrian hamsters (*Mesocricetus auratus*) of 5-6 weeks of age (average weight 60-80 grams) were purchased from Janvier Laboratories and handled under specific pathogen-free conditions. The animals were housed and manipulated in isolators in a Biosafety level-3 facility, with *ad libitum* access to water and food. Before manipulation, animals underwent an acclimation period of one week.

Animals were anesthetized with an intraperitoneal injection of 200 mg/kg ketamine (Imalgène 1000, Merial) and 10 mg/kg xylazine (Rompun, Bayer), and received one single subcutaneous injection of 200 μL of freshly-diluted ivermectin (I8898, Sigma-Aldrich) at the classical anti-parasitic dose of 400 μg/kg^32^ (or at 100-200 μg/kg for the dose-response experiment). Non-treated animals received one single subcutaneous injection of 200 μL of physiological solution. 100 μL of physiological solution containing 6×10^4^ PFU of SARS-CoV-2 was then administered intranasally to each animal (50 μL/nostril). Mock-infected animals received the physiological solution only.

Infected and mock-infected animals were housed in separate isolators and all hamsters were followed-up daily during four days at which the body weight and the clinical score were noted. The clinical score was based on a cumulative 0-4 scale: ruffled fur, slow movements, apathy, stress when manipulated.

At day 3 post-infection (dpi), animals underwent a food finding test to assess olfaction as previously described^24,33^. Briefly, 24 hours before testing, hamsters were fasted and then individually placed into a fresh cage (37 x 29 x 18 cm) with clean standard bedding for 10 minutes. Subsequently, hamsters were placed in another similar cage for 2 minutes when about 5 pieces of cereals were hidden in 1.5 cm bedding in a corner of the test cage. The tested hamsters were then placed in the opposite corner and the latency to find the food (defined as the time to locate cereals and start digging) was recorded using a chronometer. The test was carried out during a 15 min period. As soon as food was uncovered, hamsters were removed from the cage. One minute later, hamsters performed the same test but with visible chocolate cereals, positioned upon the bedding. The tests were realized in isolators in a Biosafety level-3 facility that were specially equipped for that. At 4 dpi, animals were euthanized with an excess of anesthetics (ketamine and xylazine) and exsanguination^34^, and samples of nasal turbinates and lungs were collected and immediately frozen at −80°C.

### RNA isolation and transcriptional analyses by quantitative PCR from golden hamsters’ tissues

Frozen tissues were homogenized with Trizol (15596026, Invitrogen) in Lysing Matrix D 2 mL tubes (116913100, MP Biomedicals) using the FastPrep-24™ system (MP Biomedicals) at the speed of 6.5 m/s during 1 min. Total RNA was extracted using the Direct-zol RNA MicroPrep Kit (R2062, Zymo Research: nasal turbinates) or MiniPrep Kit (R2052, Zymo Research: lung) and reverse transcribed to first strand cDNA using the SuperScript™ IV VILO™ Master Mix (11766050, Invitrogen). qPCR was performed in a final volume of 10□μL per reaction in 384-well PCR plates using a thermocycler (QuantStudio 6 Flex, Applied Biosystems). Briefly, 2.5□μL of cDNA (12.5 ng) were added to 7.5□μL of a master mix containing 5□μL of Power SYBR green mix (4367659, Applied Biosystems) and 2.5□μL of nuclease-free water with nCoV_IP2 primers (nCoV_IP2-12669Fw: 5’-ATGAGCTTAGTCCTGTTG-3’; nCoV_IP2-12759Rv: 5’-CTCCCTTTGTTGTGTTGT-3’) at a final concentration of 1□μM^35^. The amplification conditions were as follows: 95°C for 10□min, 45 cycles of 95°C for 15□s and 60°C for 1 min; followed by a melt curve, from 60□°C to 95□°C. Viral load quantification of hamster tissues was assessed by linear regression using a standard curve of eight known quantities of plasmids containing the *RdRp* sequence (ranging from 10^7^ to 10° copies). The threshold of detection was established as 200 viral copies/μg of RNA. The Golden hamster gene targets were selected for quantifying host inflammatory mediator transcripts in the tissues using the *Hprt* (hypoxanthine phosphoribosyltransferase) and the *γ-actin* genes as reference (Table 1). Variations in gene expression were calculated as the n-fold change in expression in the tissues from the infected hamsters compared with the tissues of the uninfected ones using the 2^−ΔΔCt^ method^36^.

**Table 1.**
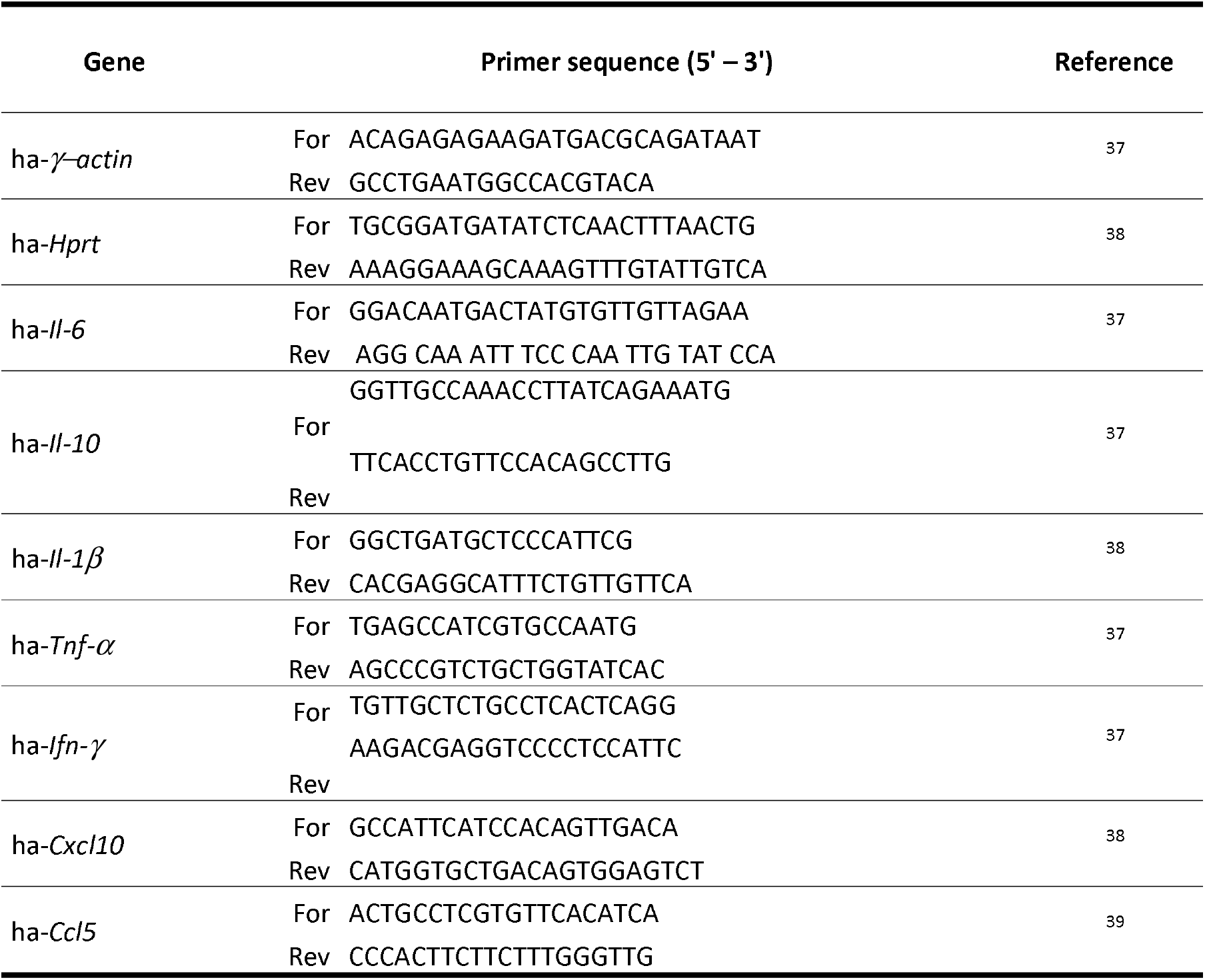
Primer sequences used for qPCR in the golden hamster tissues.

### Droplet digital PCR (ddPCR)

*Reverse transcription:* 200 ng of RNA was reverse transcribed using iScript Advanced cDNA Synthesis kit for RT-qPCR (1702537, Bio-Rad) according to the manufacturer’s specifications. *Quantitative PCR for γ-actin and Hprt reference genes:* Real-time PCR was performed in a CFX96 qPCR machine (Bio-Rad). All samples were measured in duplicate. The 10 μL PCR reaction included 0.8 ng of cDNA, 1× Powerllp PCR master mix (A25742, Applied Biosystems) and 0.5 μM of each primer (Table 1). The reactions were incubated in a 96-well optical plate at 95°C for 2 min, followed by 40 cycles of 95°C for 15s and 60°C for 1 min. *Droplet digital PCR:* ddPCR reactions were performed on the QX200 Droplet Digital PCR system according to manufacturer’s instructions (Bio-Rad). Briefly, reaction mixture consisted in 10 μL ddPCR Supermix for probe no dUTP (1863023, Bio-Rad), 0.25 to 1 ng of cDNA, primers and probes for E/IP4 and N/nsp13 duplex reactions used at concentration listed in Table 2 in a final volume of 20 μL. PCR amplification was conducted in a iCycler PCR instrument (Bio-Rad) with the following condition: 95 °C for 10 min, 40 cycles of 94 °C for 30 s with a ramping of 2°/s, 59 °C for 1 min with a ramping of 2°/s, followed by 98 °C for 5 min with a ramping of 2°/s and a hold at 4 °C. After amplification, the 96-well plate was loaded onto the QX200 droplet reader (Bio-Rad) that measures automatically the fluorescence intensity in individual droplets. Generated data were subsequently analyzed with QuantaSoft™ software (Bio-Rad) based on positive and negative droplet populations. Data are expressed as CPD (copy per droplets) normalized to *γ-actin* and *Hprt* reference genes relative expression.

**Table 2.**
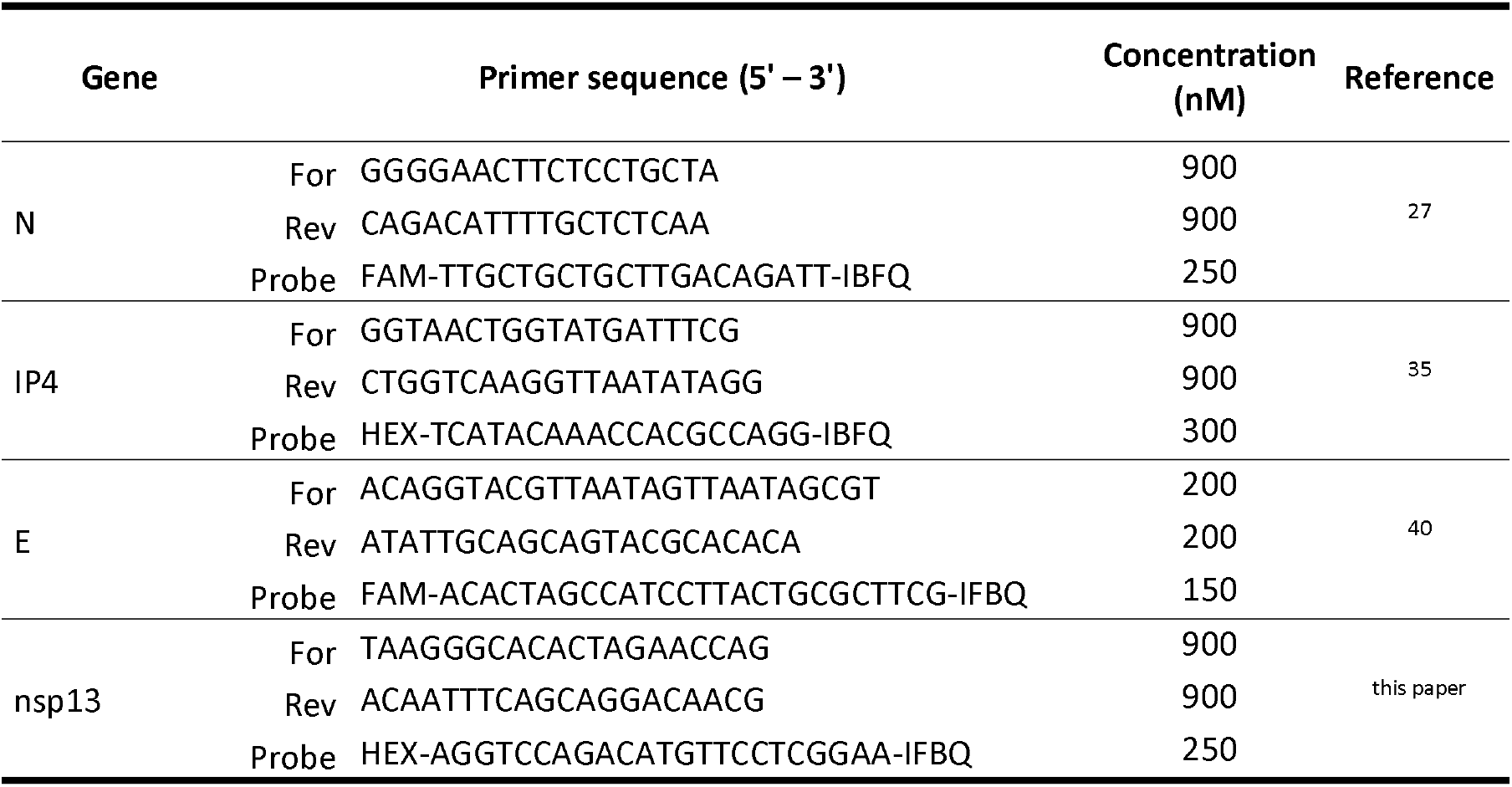
Primer and probes sequences used for ddPCR in the golden hamster lung.

### Viral titration in golden hamsters’ lung

Frozen lungs fragments were weighted and homogenized with 1 mL of ice-cold DMEM supplemented with 1% penicillin/streptomycin (15140148, Thermo Fisher) in Lysing Matrix M 2 mL tubes (116923050-CF, MP Biomedicals) using the FastPrep-24™ system (MP Biomedicals), and the following scheme: homogenization at 4.0 m/s during 20 sec, incubation at 4°C during 2 min, and new homogenization at 4.0 m/s during 20 sec. The tubes were centrifuged at 10.000 x g during 1 min at 4°C, and the supernatants were titrated on Vero-E6 cells by classical plaque assays using semisolid overlays (Avicel, RC581-NFDR080I, DuPont)^31^.

### Statistics

Statistical analysis was performed using Prism software (GraphPad, version 9.0.0, San Diego, USA), with *p* < 0.05 considered significant. Quantitative data was compared across groups using Log-rank test or two-tailed Mann-Whitney test.

## Acknowledgements

The SARS-CoV-2 strain was supplied by the National Reference Centre for Respiratory Viruses hosted by Institut Pasteur (Paris, France) and headed by Dr. Sylvie van der Werf. The human sample from which strain 2019-nCoV/IDF0372/2020 was isolated has been provided by Dr. X. Lescure and Pr. Y. Yazdanpanah from the Bichat Hospital (Paris, France). This work was supported by Institut Pasteur TASK FORCE SARS COV2 (NicoSARS project and NeuroCovid project) and received help from the European Union’s Horizon 2020 Framework Programme for Research and Innovation under Specific Grant Agreement No. 945539 (Human Brain Project SGA3). We would like to thank Marion Berard, Laeticia Breton and Rachid Chennouf for their help in implementing experiments in the Institut Pasteur animal facilities. We thank Arnaud Tarantola and Andrew Holtz for critical reading of the manuscript.

## Author Contributions

JPC and HB conceived the experimental hypothesis.

GDM, FLaz, FLar and HB designed the experiments.

GDM, FLaz, FLar, LF, LK and AM performed the experiments.

GDM, FLaz, FLar, LF, AM, PP, ML and PML analyzed the data.

GDM, JPC and HB wrote the manuscript.

## Additional information

Extended data is available for this paper.

## Reporting Summary

Further information on research design is available in the Nature Research Reporting Summary linked to this article.

## Data availability

All the data generated or analyzed during this study are included in this article along with its extended data.

---

**Extended data Fig. 1.**
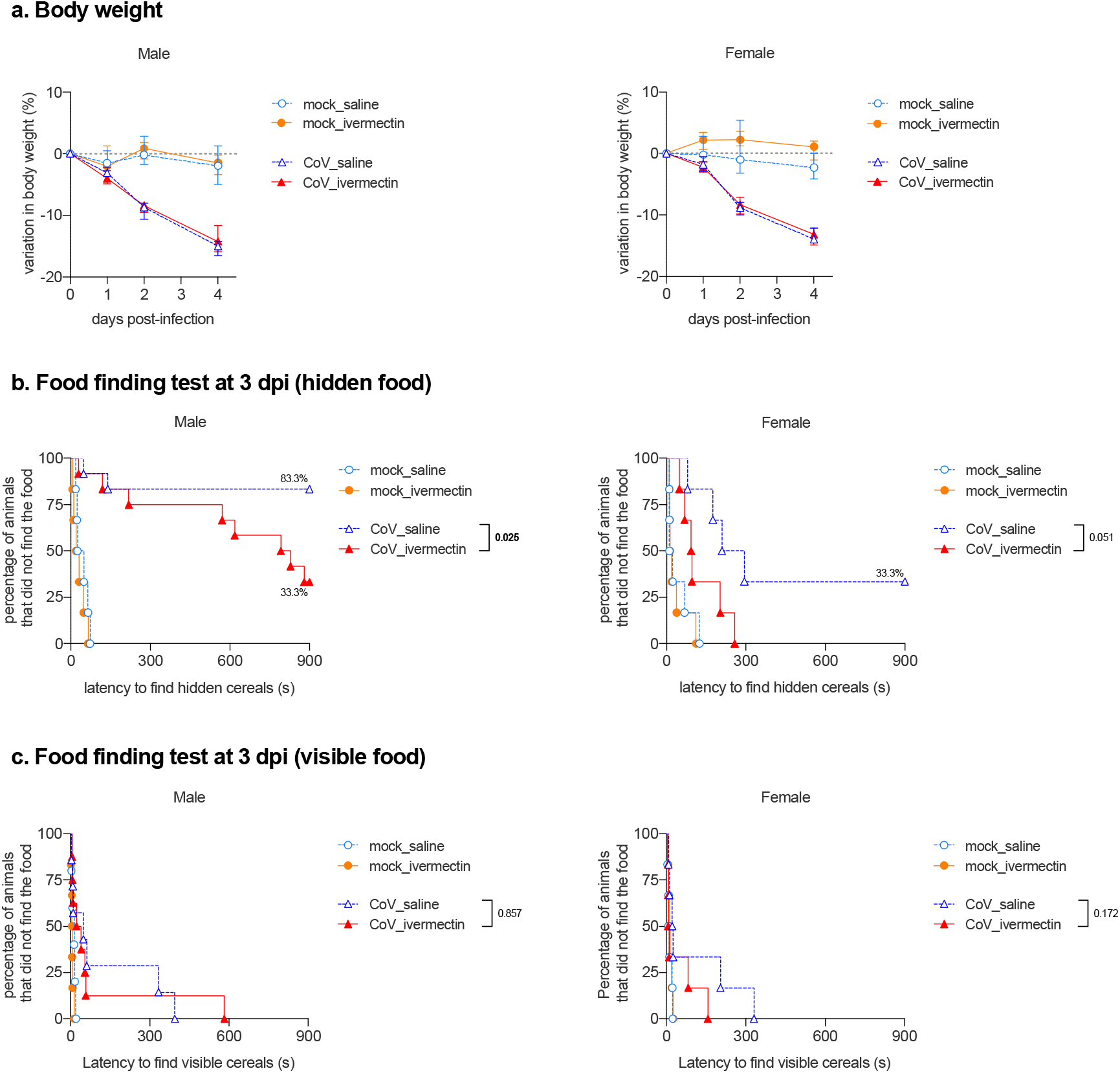
Body weight variation and Food finding test. **a,** progression of body weight in male and female hamsters, mock-infected or SARS-CoV-2 infected, treated with saline or with 400 μg/kg ivermectin. **b.** curves represent the percentage of animals that did not find the hidden (buried) food. **c.** curves represent the percentage of animals that did not find the visible (unburied) food. Food finding assays were performed at 3 days post-infection, n=12/group (males CoV_saline and males CoV_i verm ectin). n=6/group (females CoV_saline and females CoV-ivermectin). n=4/group (males and females mock_ivermectin). n=3/group (males and females mock_saline). Log-rank (Mantel-Cox) test. The p value is indicated in bold when significant at a 0.05 threshold. Data were obtained from three independent experiments for males and two independent experiments for females.

**Extended data Fig. 2.**
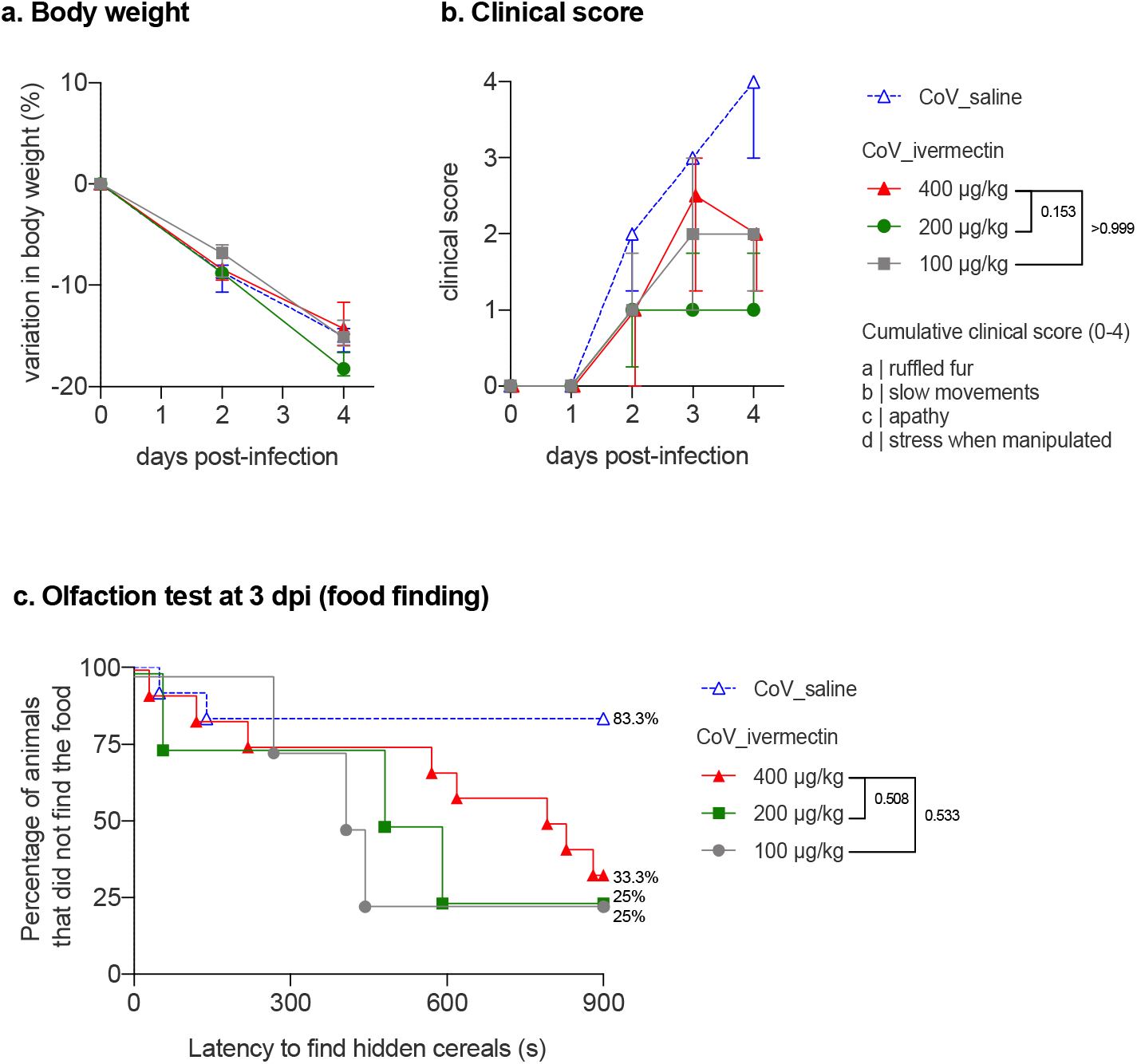
Clinical aspects of SARS-CoV-2-infected male hamsters and treated with different doses of ivermectin. **a**, progression of body weight in male hamsters, treated with saline or with 400 μg/kg, 200 μg/kg or 100 μg/kg ivermectin. **b**, clinical score based on a cumulative 0-4 scale: ruffled fur; slow movements; apathy; stress when manipulated. **c**, olfaction deficit based on the buried food finding test. Curves represent the percentage of animals that did not find the buried food. Food finding assays were performed at 3 days post-infection. n=12/group (CoV_saline and CoV_ivermectin 400 μg/kg, as shown in Fig. 1) or n=4 (CoV_ivermectin 200 μg/kg and 100 μg/kg). Mann-Whitney test at 4 dpi (b) and Log-rank (Matel-Cox) test (c). The p value is indicated in bold when significant at a 0.05 threshold. Symbols indicate the median ± interquartile range.

**Extended data Fig. 3.**
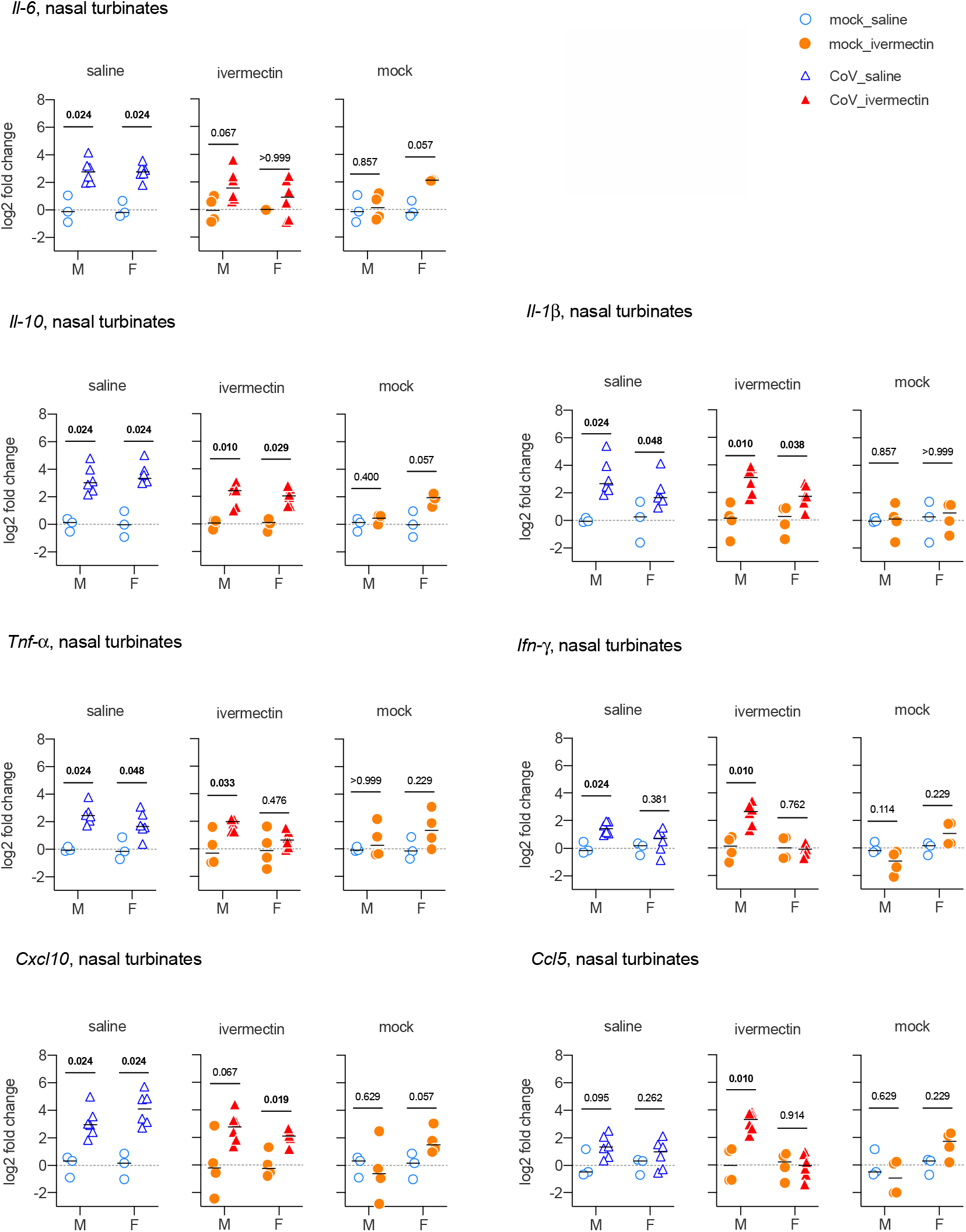
Immunological aspects in the nasal turbinates at 4 days post-infection of SARS-CoV-2-infected hamsters with and without ivermectin treatment. Cytokines and chemokines transcripts in the nasal turbinates at 4 dpi. n=6/group (CoV_saline and CoV-ivermectin), n=4/group (mock_ivermectin), n=3/group (mock_saline). Mann-Whitney test. The p value is indicated in bold when significant at a 0.05 threshold. Horizontal lines indicate medians. M: male hamsters; F: female hamsters. Data were obtained from two independent experiments for each sex.

**Extended data Fig. 4.**
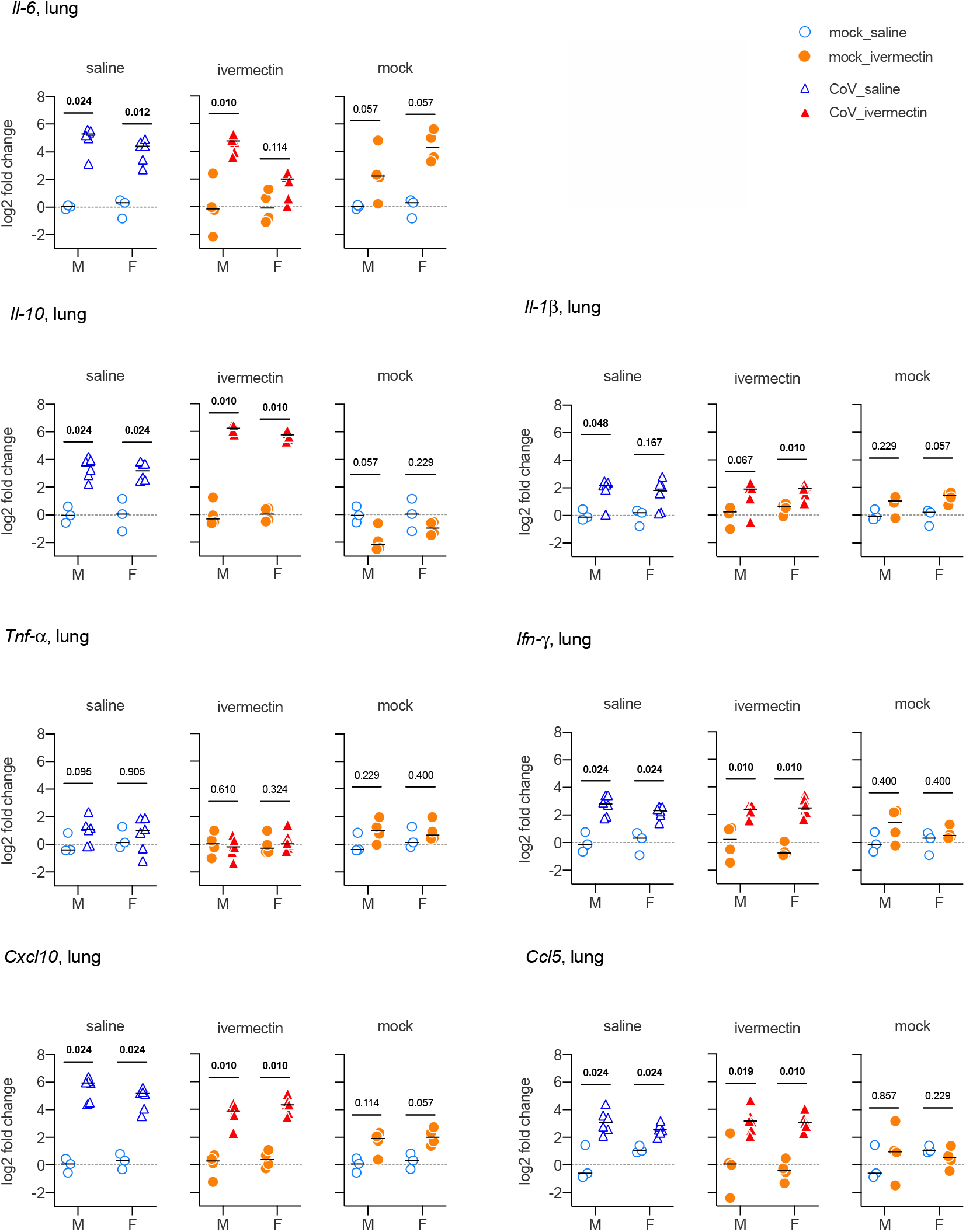
Immunological aspects in the lungs at 4 days post-infection of SARS-CoV-2-infected hamsters with and without ivermectin treatment. Cytokines and chemokines transcripts in the lungs at 4 dpi. n=6/group (CoV_saline and CoV-ivermectin), n=4/group (mock_ivermectin), n=3/group (mock_saline). Mann-Whitney test. The p value is indicated in bold when significant at a 0.05 threshold. Horizontal lines indicate medians. M: male hamsters; F: female hamsters. Data were obtained from two independent experiments for each sex.

**Extended data Fig. 5.**
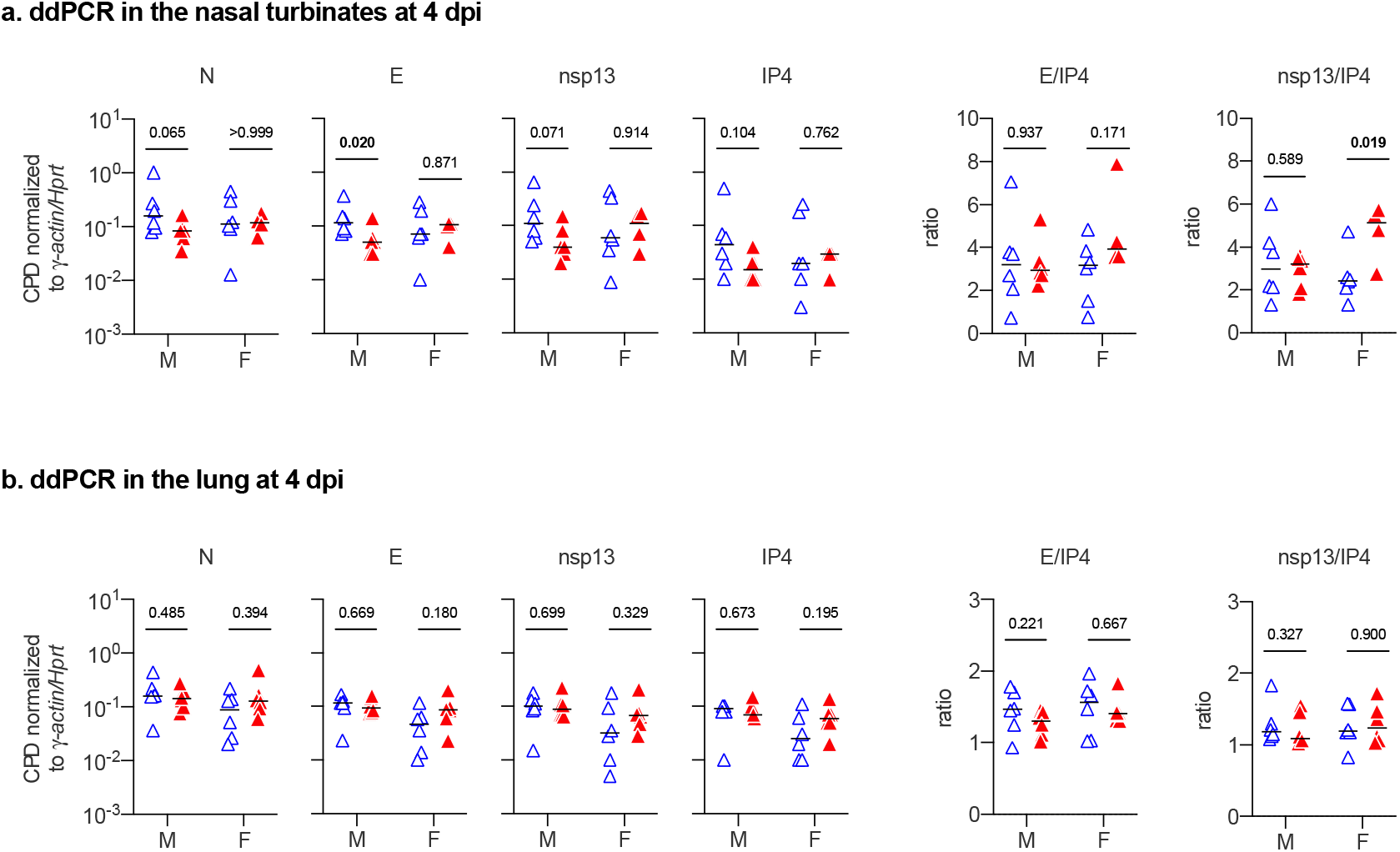
Virologic aspects in the nasal turbinates and in the lungs at 4 days post-infection of SARS-CoV-2-infected hamsters with and without ivermectin treatment. **ab**, viral gene expression of N (nucleocapsid), E (envelope), nsp13 (non-structural protein 13), IP4 (RdRp, RNA-dependent RNA polymerase), and E/IP4 and nsp13/IP4 ratios determined by digital droplet PCR (ddPCR) in the nasal turbinates (a) and in the lungs (b) at 4 dpi. Data are expressed as CPD (copy per droplets) normalized to *γ-actin* and *Hprt* reference genes relative expression. n=6/group (except nasal turbinates from female CoV_ivermectin, where n=4). Mann-Whitney test. The p value is indicated in bold when significant at a 0.05 threshold. Horizontal lines indicate the medians. M: male hamsters; F: female hamsters. Data were obtained from two independent experiments for each sex.

